# Bioengineered model of the human motor unit with physiologically functional neuromuscular junctions

**DOI:** 10.1101/2020.05.04.076778

**Authors:** Rowan P. Rimington, Jacob W. Fleming, Andrew J. Capel, Patrick C. Wheeler, Mark P. Lewis

## Abstract

Investigations of the human neuromuscular junction (NMJ) have predominately utilised experimental animals and model organisms. Consequently, there remains a paucity of data regarding the development of the human NMJ and a lack of systems that enable temporal investigation of the motor unit. This work addresses this need, providing the methodologies to bioengineer 3D models of the human motor unit. Separate maturation of primary human skeletal muscle and iPSC derived motor neurons seeks to accurately represent neuromuscular development via controlled addition of motor axons following primary myogenesis. Spheroid cultures of motor neuron progenitors augmented the transcription of OLIG2, ISLET1 and SMI32 motor neuron mRNAs ∼400, ∼150 and ∼200-fold respectively. Axon projections of adhered motor neuron spheroids exceeded 1000μm in monolayer, with transcription of SMI32 and VACHT mRNAs further enhanced in a concentration dependent manner within optimised 3D type I collagen extracellular matrices. Bioengineered skeletal muscles produce functional forces, demonstrate increased acetylcholine receptor (AChR) clustering, and transcription of MUSK and LRP4 mRNAs indicating enhanced organisation of the post-synaptic membrane. Dosed integration of motor neuron spheroids outlined the motor pool required to functionally innervate muscle tissues in 3D, generating physiologically functional human NMJs that evidence pre- and post-synaptic membrane and motor nerve terminal co-localisation. Spontaneous firing was significantly elevated in 3D motor units, confirmed to be driven by the motor nerve via antagonistic inhibition of the AChR. Finally, functional analyses outlined decreased time to peak twitch and half relaxation times, indicating enhanced physiology of excitation contraction coupling of NMJs within innervated motor units.

## 1. Introduction

The neuromuscular junction (NMJ) is the specialised synapse between the post-synaptic skeletal muscle fibre and pre-synaptic terminal of the efferent motor nerve axon. Its function is to elicit contraction of the peripheral musculature and consequently control locomotion of the skeleton via a co-ordinated mechanosensory-motor circuit. The NMJ comprises an organised assembly of protein complexes (Agrin-Lrp4-MuSK-Dok7) that form the highly specialised synaptic basal lamina; the function of which is to stabilise the synapse and facilitate the intricate biochemical processes that initiate muscular excitability and contraction.^1,2^ Neuronal action potential induces an influx of calcium ions at the motor nerve terminal that elicits fusion of synaptic vesicles at active zones on the terminal membrane. Vesicle fusion mediates the release of the neurotransmitter acetylcholine (ACh) which diffuses through the synaptic cleft and binds nicotinic acetylcholine receptors (AChR) on the post-synaptic membrane.^3^ This depolarisation opens voltage-gated sodium ion channels and generates the action potential required for excitation contraction coupling and muscular contraction.^4^

To date, study of neuromuscular development and physiological function has predominantly utilised experimental rodent, zebrafish and drosophila models.^5–7^ As the NMJ is implicated in a variety of neuromuscular/neurodegenerative diseases, multiple transgenic rodent models have also been produced to investigate disease specific pathology.^8,9^ Although significant advancements have been made utilising animal models, issues remain when translating animal data to human physiology.^10^ This is re-enforced by reported differences in the cellular and molecular composition of the animal NMJ compared to that of humans.^11^ Consequently, there remains a paucity of data specific to the development and function of the human NMJ, and a requirement to temporally analyse development, disease or responses to external stimuli.

Human cellular systems that enable the dynamic modelling of the NMJ have been significantly advanced by the ability to generate human motor neurons from induced pluripotent stem cells (iPSCs).^12,13^ Several systems and methodologies have now been reported that combine human skeletal muscle and iPSC motor neurons in monolayer, however, these two-dimensional (2D) systems lack the physiological complexity of the NMJ.^14–18^ Three-dimensional (3D) co-culture systems enable the advanced modelling of the neuromuscular system due to incorporation of extracellular matrix proteins and mechanical forces that enable enhanced biological development.^19^ Further, 3D tissue systems provide a flexibility in manipulation (genetic modification/exogenous treatments), throughput, and rapid experimental iteration that cannot be achieved in animal models. 3D microfluidic systems using C2C12 skeletal myoblasts innervated with iPSC motor neurons,^20^ and classical engineered tissues using primary rat derived muscle and motor neurons have been reported.^21,22^ However, neither system fully utilises human cellular materials and consequently cannot replicate human NMJ physiology.

Only two successful methodologies to functionally innervate human skeletal muscle with spinal motor neurons in 3D have been reported to date.^23,24^ The first example of human NMJ 3D culture was produced by Osaki et al, with the microfluidic device enabling the separate maturation of iPSC derived motor neurons in spheroids and iPSC muscle tissue. Although excellent functional evaluation was performed, limited biochemical characterisation of NMJ formation was undertaken. Further, while using iPSC myogenic precursors is ideal for autologous disease modelling, it removes the endogenous supporting populations evident within biopsied primary human skeletal muscle that contribute to its physiology and function. Afshar Bakooshli et al reported the first engineered tissue with primary human skeletal muscle and iPSC motor neurons comprising the most sophisticated functional and biochemical characterisation to date. Despite this advance, iPSC motor neurons were harvested from monolayer populations and combined with myogenic precursors in matrices specifically tailored for skeletal muscle at experimental onset. To maximise neuronal maturity and accurately model human NMJ development, 3D cultures, optimised cell-specific extracellular matrices and methods that enable precision in the time of neuronal addition dependent on the stage of myogenesis are required. Further, while previous work has focussed on disease, no human model has evidenced functional changes indicative of enhanced muscular excitability for the study of physiological neuromuscular contraction.

Here, we present methodologies to produce 3D tissues of the human neuromuscular motor unit with physiologically functional NMJs, using primary skeletal muscle and iPSC derived motor neurons. Producing 3D spheroid cultures of motor neurons accelerates development and enhances transcription of mature SMI32 mRNAs 200-fold compared to monolayer equivalents, extending axons >1000μm in length. Optimising type I collagen matrices further enhances motor neuron spheroid maturation 6-fold and provides a loading mechanism for delivery to primary human skeletal muscle tissue within a freely available, open-source 3D printed system.^25^ The separate maturation of both neuronal and myogenic populations allows for control of motor neuron addition to engineered muscle depending on stage of developmental myogenesis. Further, this is the first research to evidence enhanced physiology of time to peak contraction and half relaxation time, following innervation in engineered tissues.

## 2. Results

### 2.1. 3D spheroid culture of iPSC derived motor neuron progenitors accelerates development and enhances maturation

To enhance maturity of iPSC motor neurons, progenitor cells were combined within microplates to form developmental spheroids and compared against progenitors cultured in monolayer. iPSC derived motor neuron progenitor spheroids, positive for Olig-2, HB-9, SMI-32 and cytoskeletal marker Tuj-1 enabled continual development across a 3 week culture period (Fig.1a). In comparison, monolayer cultures also developed positivity of mature marker SMI-32, however, by week 3 adherent cells were observed to be losing phenotype and adherence to coated culture surfaces (Fig.1b). Transcriptional analysis of motor neuron lineage outlined 20-fold increases in developmental regulator PAX6 mRNA after 2 days spheroid culture (P ≤ 0.001), decreasing thereafter. Contrastingly, monolayer progenitor PAX6 mRNA expression increased from day 2 to day 7 and maintained this elevated level throughout 3 weeks culture, significantly compared to 3D spheroids after 21 days (P ≤ 0.05, Fig.1c). Expression of progenitor marker OLIG2 mRNA was significantly elevated in spheroid cultures after 2 (P ≤ 0.01) and 7 (P ≤ 0.01) days, peaking at 400-fold after 14 days culture (P ≤ 0.001) prior to decreasing below that of monolayer cells after 21 days (P ≤ 0.01, Fig.1d). This transcriptional pattern was also evident in motor neuron maker ISLET1 mRNA being significantly elevated in spheroid cultures after 2 (P ≤ 0.001), 7 (P ≤ 0.001) and 14 days (P ≤ 0.05) culture (Fig.1e). mRNA of mature motor neuron marker SMI32 was also significantly elevated after 2 days (P ≤ 0.001) in spheroid culture, however continued to increase in transcription throughout development after 14 (P ≤ 0.001) and 21 days (P ≤ 0.001, Fig.1f), 200- fold compared to monolayer. Proliferative maker MKI67 mRNA was elevated after 2 days in motor neuron spheroids (P ≤ 0.05), however this response was then decreased after 7 (P ≤ 0.05), 14 (P ≤ 0.01) and 21 days culture (P ≤ 0.01), with monolayer populations demonstrating increased proliferation throughout development (Fig.1g).

**Fig.1.**
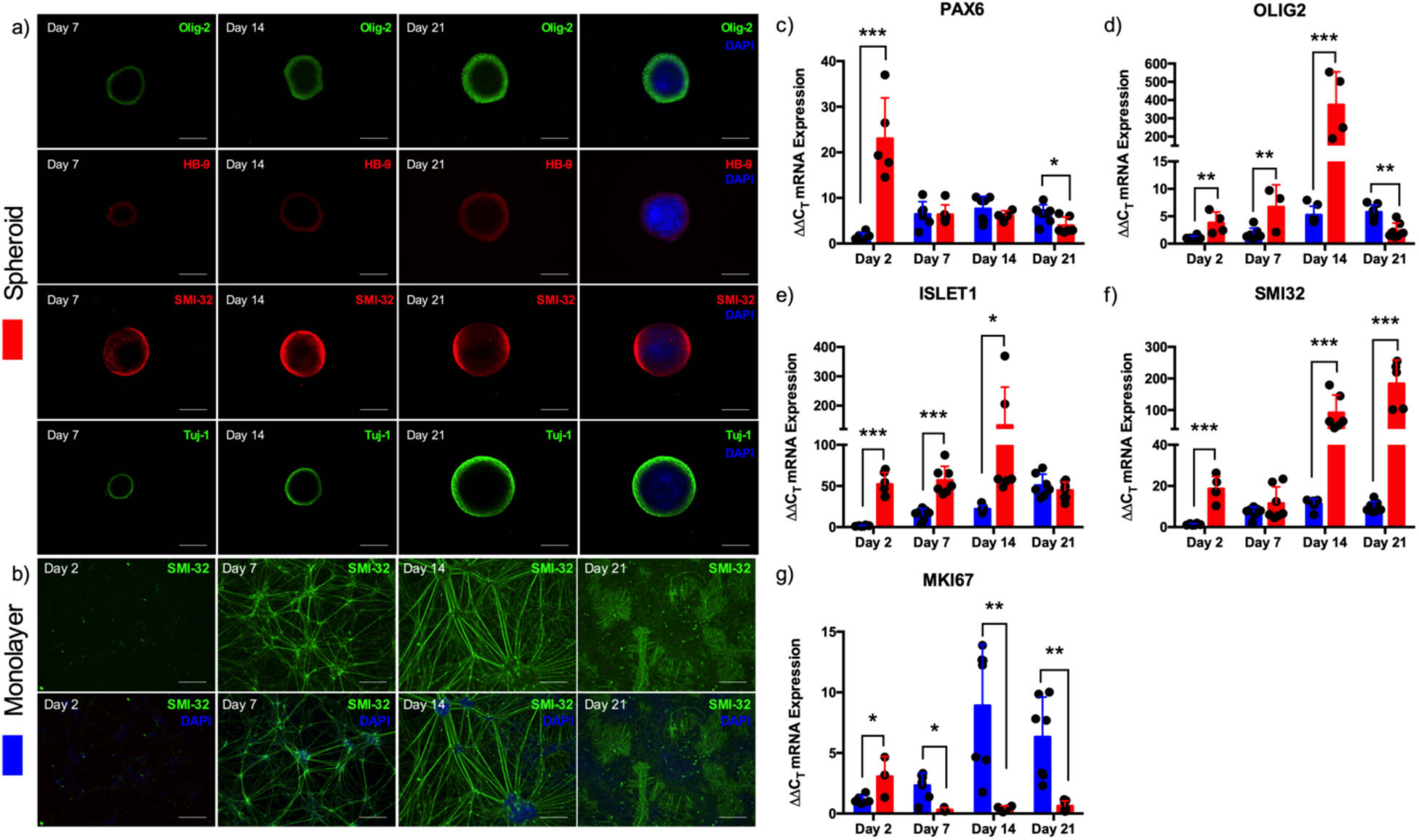
Spheroid culture of iPSC derived motor neuron progenitors accelerates transcription of developmental and mature mRNAs. (a) Motor neuron progenitors cultured as 3D spheroids for 7, 14 and 21 days positive for Olig-2, HB-9, SMI-32 and Tuj-1. (b) Motor neuron progenitor cells cultured in monolayer for 2, 7, 14 and 21 days positive for SMI-32. (c) mRNA gene expression for neuronal stem cell marker PAX6, (d) progenitor marker OLIG2, (e) motor neuron marker ISLET1, (f) mature motor neuron marker SMI32 and (g) proliferative marker MKI67. Data presented ± standard deviation (SD). *P ≤ 0.05, **P ≤ 0.01, and ***P ≤ 0.001. Scale bars = 200µm.

### 2.2. iPSC motor neuron spheroids axonal extension is dependent on developmental stage

To investigate the axonal extension potential of motor neurons at different developmental stages, spheroids were removed after 7, 14, and 21 days differentiation, adhered to culture well plates and left for 5 days (Fig.2a). Spheroids extended significantly longer axons after 14 days development, compared to 7 (P ≤ 0.01) and 21 days (Fig.2b, P ≤ 0.001) indicating that extension of Tuj-1+ axons are temporally regulated in development (P ≤ 0.001). Maximum lengths of axons at all three developmental stages typically exceeded 1000µm, with axons significantly longer after 14 days compared to 21 (P ≤ 0.01). SMI-32 positivity of spheroid axons remains consistent (>80%) at all stages of development (P ≥ 0.05 Fig.2d), indicating comparable high levels of axon maturity. Considering previous mRNA expression with axon extension data, it was determined that 14 days spheroid culture was optimal for motor neuron development before adherent culture. However, both earlier and later time-points produced viable motor neurons of enhanced development extending long axons.

**Fig.2.**
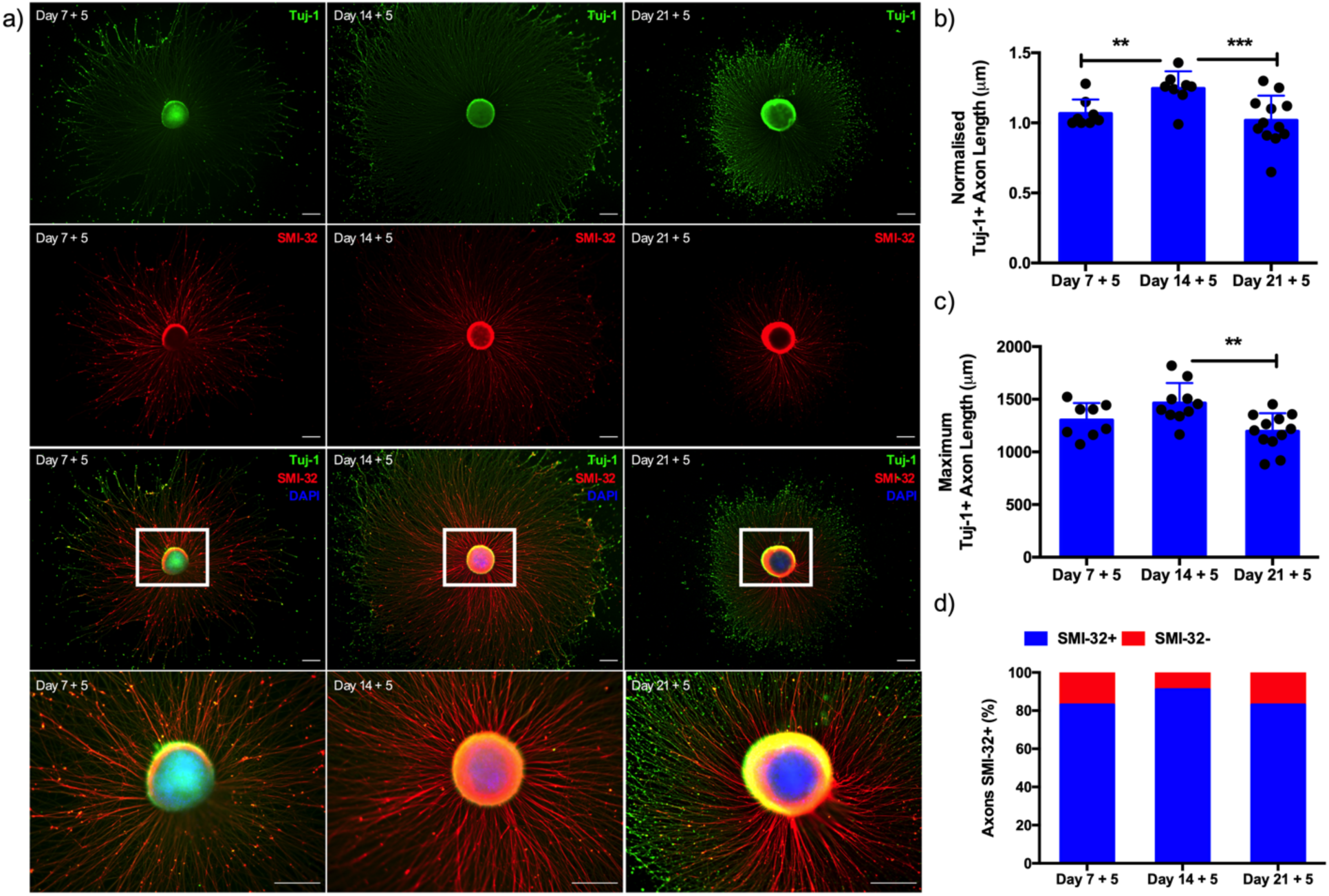
iPSC motor neuron spheroids axonal extension potential is determined via developmental stage. (a) Motor neuron progenitors cultured as 3D spheroids at different stages of development (7, 14 and 21 days) adhered in monolayer culture extend axons over 5 days in culture. (b) Normalised Tuj-1 positive (+) axon length after 5 days adherent culture following seeding at days 7, 14 and 21 days of maturation. (c) Maximum Tuj-1 axon lengths (µm) at each stage of maturation. (d) Percentage (%) axons positive for mature heavy neurofilament SMI-32 after 5 days adherent culture at different development stages. Data indicative of minimum n=8 per time- point and presented ± standard deviation (SD). **P ≤ 0.01, and ***P ≤ 0.001. Scale bars = 200µm.

### 2.3. Culture of iPSC motor neuron spheroids within 3D collagen I matrices exhibit concentration dependent enhancement in maturity

To mimic 3D *in vivo* environments, motor neuron spheroids were cultured to maturity (14 days), resuspended within 3D type I collagen matrices in a concentration dependent manner (Fig.3a), and again cultured for a further 5 days. Spheroids were then analysed for mRNA markers of maturity (SMI32), vesicular transporter of acetylcholine (VACHT) and microtubule-associated protein tau (MAPT); indicative of axon extension, compared with 14 day monolayer spheroids (Fig.2). Suspension within 3D collagen environments further enhanced SMI32 mRNA expression, with transcription of this mature gene elevated 6-fold in 1mg/mL hydrogel compositions compared to monolayer (P ≤ 0.001, Fig.3b). This trend was also evident for the VACHT mRNA, with addition to the 3D environment increasing expression, specifically in 1mg/mL type I collagen matrices compared to monolayer spheroids (Fig.3c, P ≤ 0.05). Although not statistically significant, 50% increases in MAPT mRNA were also observed in 1mg/mL conditions compared to monolayer (Fig.3d). Consequently, 1mg/mL concentrations of type I collagen were selected as the ideal loading matrix for iPSC motor neuron spheroids in this work.

**Fig.3.**
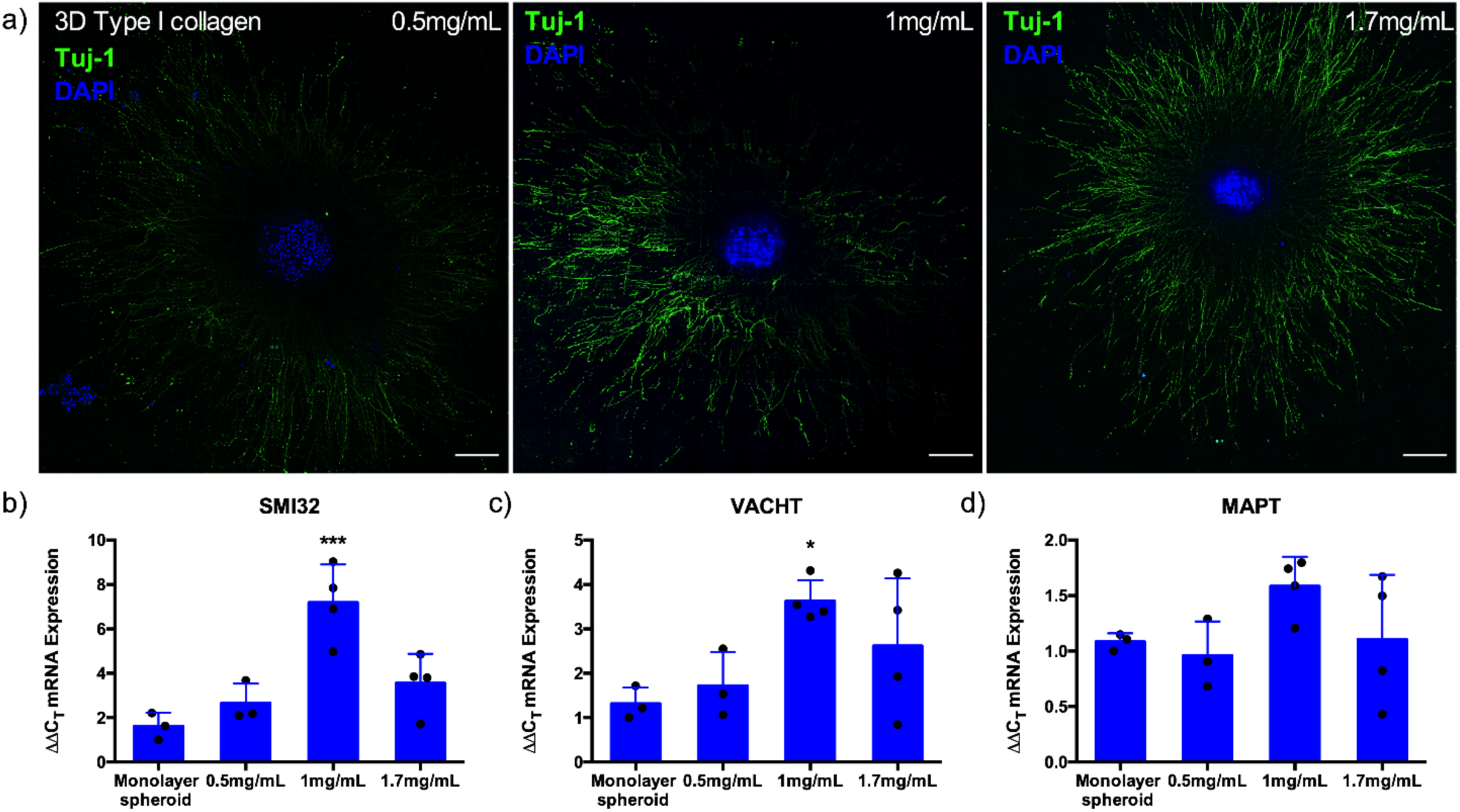
Culture of iPSC motor neuron spheroids within 3D collagen I matrices exhibit concentration dependent enhancement in mRNA expression of heavy neurofilament (SMI-32), vesicular transporter of acetylcholine (VAChT) and axonal extension (MAPT). (a) Confocal tile scans of motor neuron progenitor spheroids suspended within 3D collagen I matrices of different concentrations (0.5, 1 and 1.7mg/mL) adhere and extend axons over 5 days. (b) mRNA expression of mature motor neuron markers SMI32 and (c) VACHT, and (d) MAPT as indication of amount of axonal extension compared to adhered monolayer spheroids. Data presented ± standard deviation (SD). *P ≤ 0.05 and ***P ≤ 0.001. Scale bars = 200µm.

### 2.4. Functional 3D bioengineered primary human skeletal muscle enhances acetylcholine receptor clustering and expression of pre-synaptic mRNAs

To determine the myogenic development, differentiation of HDMCs was performed in both monolayer and 3D cultures across a 3 week period to ascertain morphological (Fig.4a) and functional (Fig.4j and k) capacity. Further, analysis of myogenic and synaptic mRNA expression, and post-synaptic receptor development were also performed to indicate the optimal window for introduction of the motor nerve input within neuromuscular co-cultures. Differences in myotube diameter were observed between monolayer and 3D tissue, and across the 3 week culture period (P *≤* 0.001). Monolayer cell cultures increased in diameter until week 2 before then reducing in width at week 3, opposed to 3D tissues that demonstrated gradual increases throughout development (Fig.4b). Linear increases in the number of AChR clusters were evident in both culture modalities, however, these were significantly elevated in 3D tissues at all time-points (P *≤* 0.001, Fig.4c). Although this increase was not reflected in the size of AChRs (P *≥* 0.05, Fig.4c), those evident in 3D tissues formed clusters with greater lacunarity as seen *in vivo* (Fig.4i). Transcriptional profiles of MYOD mRNA were consistent across both monolayer and 3D tissues, although elevations in expression were observed after 7 days in monolayer conditions (P *≤* 0.01, Fig.4e). Expression of MYOG mRNA was significantly elevated after 2 days culture in 3D tissues (P *≤* 0.001), however this decreased compared to monolayer after 1 week culture (P *≤* 0.01, Fi.g4f). Increases in MUSK mRNA in 3D tissues was apparent compared to monolayer across the entire culture period (Fig.4g), whereas LRP4 mRNA was significantly elevated immediately after 2 days (P *≤* 0.001) and again after 14 days culture in 3D tissues (P *≤* 0.05, Fig.4h). Functional readouts of 3D tissues in this work became evident after 2 weeks, with slight increases in force output evident following a further weeks culture, however, these changes were not statistically significant (P *≥* 0.05, Fig.4j, k).

**Fig.4.**
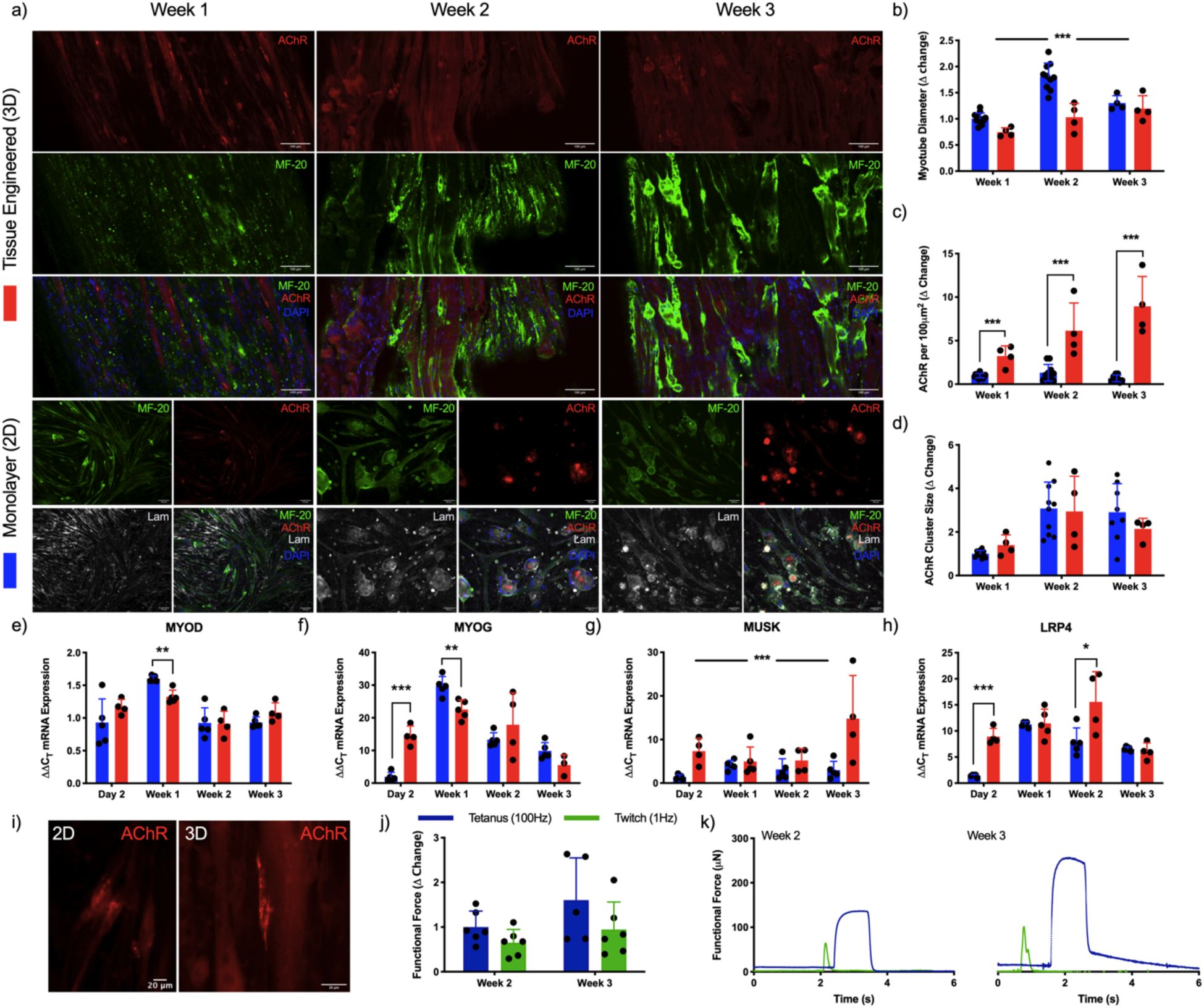
3D culture of primary human skeletal muscle enhances acetylcholine receptor clustering, produces functional force and upregulates transcription of pre-synaptic mRNAs. (a) Confocal tile scans of single z-plane (Tissue engineered 3D) and fluorescence images (Monolayer 2D) of human skeletal muscle labelled for AChR on the post-synaptic membrane, pan-myosin heavy chain (MF- 20), laminin (monolayer only) and nuclear DNA (DAPI) across 1, 2 and 3 weeks culture. Morphological analysis of (b) myotube diameter, (c) AChR density and (d) cluster size. Transcriptional analysis of myogenic; (e) MYOD and (f) MYOG, and pre-synaptic; (g) MUSK and (h) LRP4 mRNAs. (i) Zoom images of AChR in monolayer (2D) and tissue engineered (3D) skeletal muscle. (j) Functional tetanus and twitch force data and (k) representative traces in engineered tissues. Δ change indicative of normalised data. Data presented ± standard deviation (SD). * P ≤ 0.05, **P ≤ 0.01 and ***P ≤ 0.001. Scale bars = (a) 100µm, (i) 20µm.

### 2.5. Loading density of iPSC motor neuron spheroids determines functionality of neuromuscular tissues

Motor neuron spheroids were matured for 14 days as outlined (Fig.5a, i) and dosed around skeletal muscle tissues at a loading density of 2, 4 or 6 spheroids/tissue after 14 days maturation (Fig.5a, ii). Following further 2 week co-culture periods, assessment of skeletal muscle only (SkM -MN) and neuromuscular tissue force production via electrical field stimulation outlined no statistically significant increases in force when motor neurons were added following 2 weeks of myogenic differentiation (Fig.5b, d, P *≥* 0.05). Trends in functional force were, however, aligned with spontaneous contractions indicative of motor nerve input that were significantly elevated when 2 (P *≤* 0.001) and 4 MN spheroids (P *≤* 0.01) were loaded in surrounding type I collagen perimysium compared to SkM -MN controls (Fig.5c, e).

**Fig.5.**
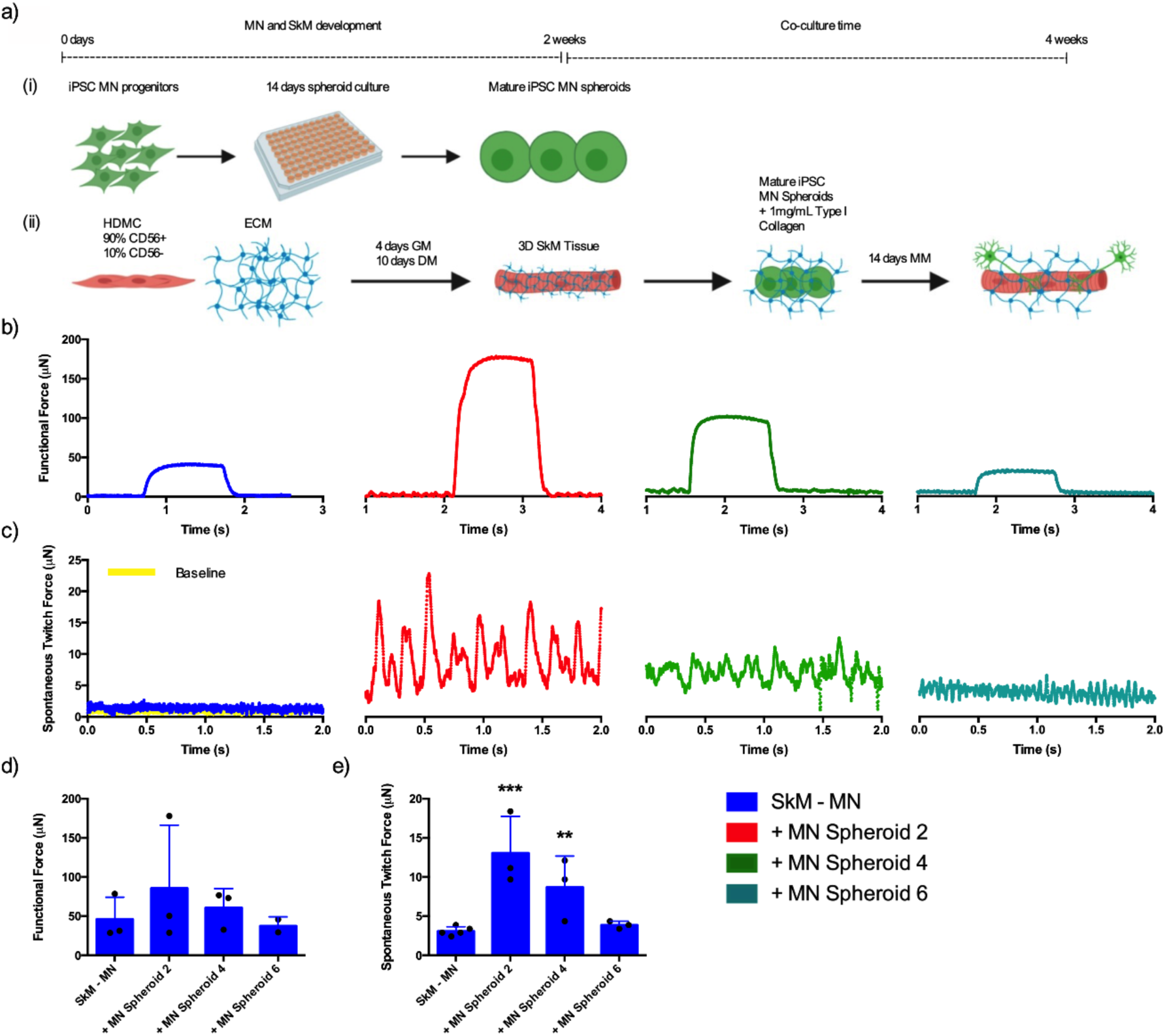
Number of iPSC motor neuron spheroids loaded in tissue engineered skeletal muscle determines functionality of neuromuscular tissues. (a) Schematic detailing method of iPSC motor neuron loading to generate neuromuscular tissues at different stages of myogenic development. (i) MN progenitors are loaded to spheroid microplates and matured for 14 days. (ii) Primary HDMCs are seeded in established type I collagen matrix, loaded into 3D printed mould and cultured to maturity for 2 weeks. Matured MNs are then added within optimised type I collagen hydrogel, seeded around SkM and cultured for a further 2 weeks. MN; Motor neuron, SkM; Skeletal muscle, HDMC; Human derived muscle cells, GM; Growth medium, DM; Differentiation medium, ECM; Extracellular matrix, MM; Motor neuron maintenance medium. (b) Addition of iPSC motor neurons with type I collagen perimysium at loading densities of 2, 4 or 6 spheroids per tissue evidence tetanic force and (c) dose dependent spontaneous twitch profiles compared to skeletal muscle only tissues. (d) Quantification of functional tetanic and (e) spontaneous twitch force output with 2, 4 or 6 motor neuron spheroids. Individual functional data points indicative of n=3 contraction profiles, totalling minimum of n=9 and maximum of n=15 per condition and presented ± standard deviation (SD). **P ≤ 0.01 and ***P ≤ 0.001.

### 2.6. Addition of iPSC derived motor neuron spheroids to matured bioengineered skeletal muscle produces functional human neuromuscular junctions

Confocal tile scans of neuromuscular tissues confirmed innervation indicated by co-localisation of the pre- (synaptic vesicle protein 2 (SV-2)) and post-synaptic (AChR) markers, and motor nerve terminal (Tuj-1, Fig.6a, e). Neuronal axons tracking along myofibers positive for myosin heavy chain and colocalising with AChR clusters (Fig.6b) were also evident, in addition to presenting classical pretzel-like morphologies (Fig.6e, f). Functionality of skeletal muscle and synaptic contacts were assessed via electrical field stimulation and analysis of spontaneously occurring twitch profiles respectively (Fig.7a). No significant differences were observed in tetanic force despite addition of the motor neurons (P *≥* 0.05), however, significant increases in spontaneous twitch force was evident (P *≤* 0.001, Fig.7b). Muscular tissue devoid of the motor nerve displayed spontaneous contractions comparable to baseline readings, which was increased 4-fold above baseline in neuromuscular tissues. To fully determine functionality between pre- and post-synaptic membranes, the AChR antagonist d-tubocurarine was added to spontaneously contracting tissues (Fig.7c). Following addition, total twitches and frequency of twitch profiles were ablated indicating inhibition of the AChR and confirming motor nerve induced contractions in neuromuscular tissues. Comparable response in maximal twitch profiles to tetanic profiles were apparent with no differences upon functional innervation (P *≥* 0.05, Fig.7d). However, measures of skeletal muscle excitability; time to peak twitch (P *≤* 0.001, Fig.7e), and relaxation; half relaxation time (P *≤* 0.01, Fig.7f), were significantly decreased following innervation (Fig.7g).

**Fig.6.**
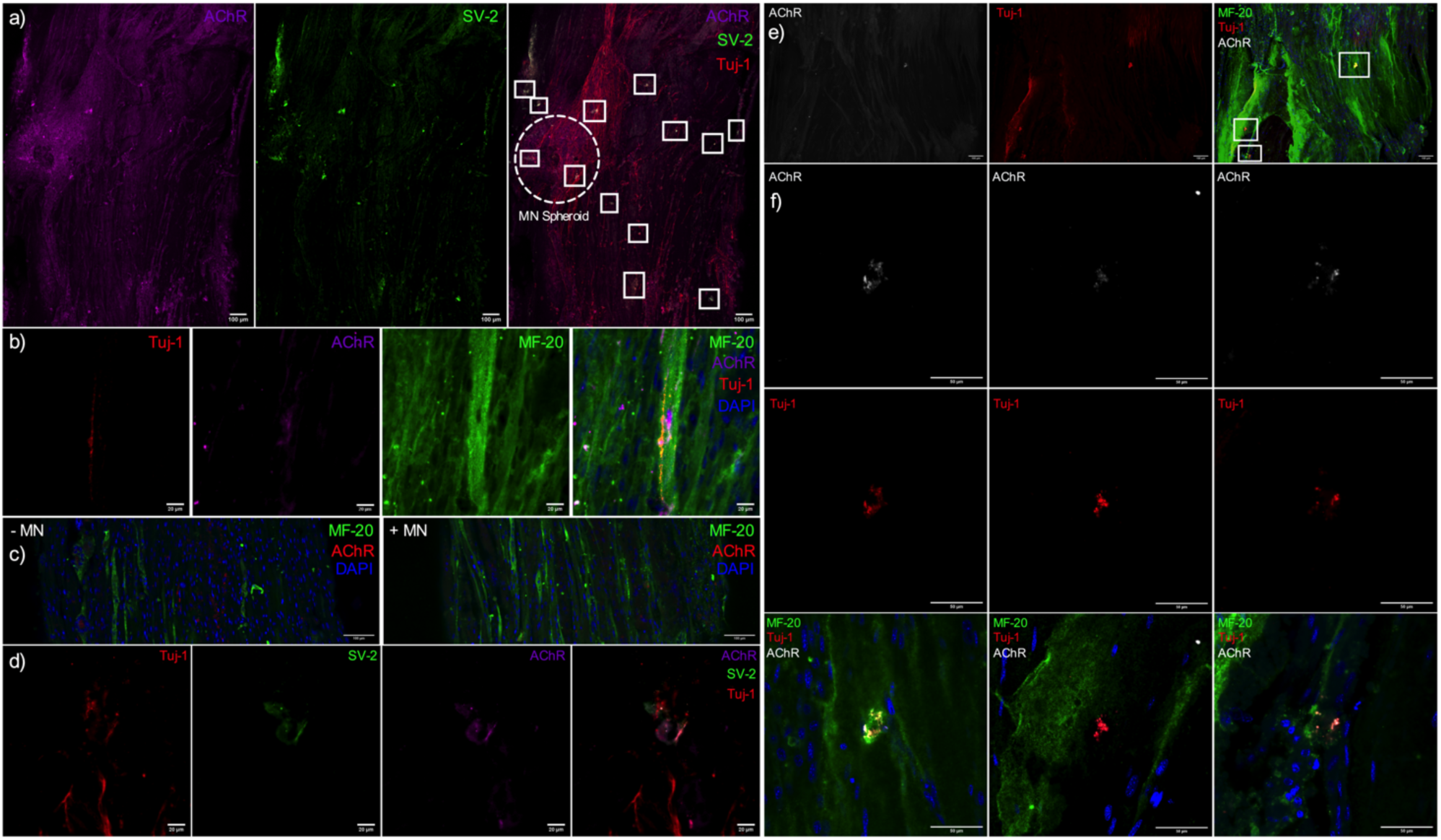
Co-localisation of pre- and post-synaptic membrane proteins, motor nerve terminals and myosin heavy chain positive fibres indicate neuromuscular junction formation in engineered tissues. (a) Confocal tile scan of neuromuscular tissue detailing AChR (left) and SV-2 (middle), with overlay (right) evidencing multiple synaptic contacts (outlined – white boxes) via co-localisation of pre- and post-synaptic markers and Tuj-1+ neuronal axons. (b) Single snap of Tuj-1 positive axon tracking down a muscle fibre positive for myosin heavy chain (MF-20), with overlapping nerve terminal and post-synaptic AChR. (c) Confocal tile scans of muscle only (-MN) and neuromuscular tissues (+MN) demonstrating comparable morphological myogenesis. (d) Zoom of individual synaptic contact as outlined in (a). (e) Confocal tile scan of neuromuscular tissues evidencing AChR, Tuj-1+ nerve terminal and myosin heavy chain co-localisation. (f) Zoom of synaptic contacts highlighted in (e) via white box outline. Images were collected across n=8 neuromuscular tissues. Scale bars = (a, c, e) 100µm, (b, d) 20µm, (f) 50µm.

**Fig.7.**
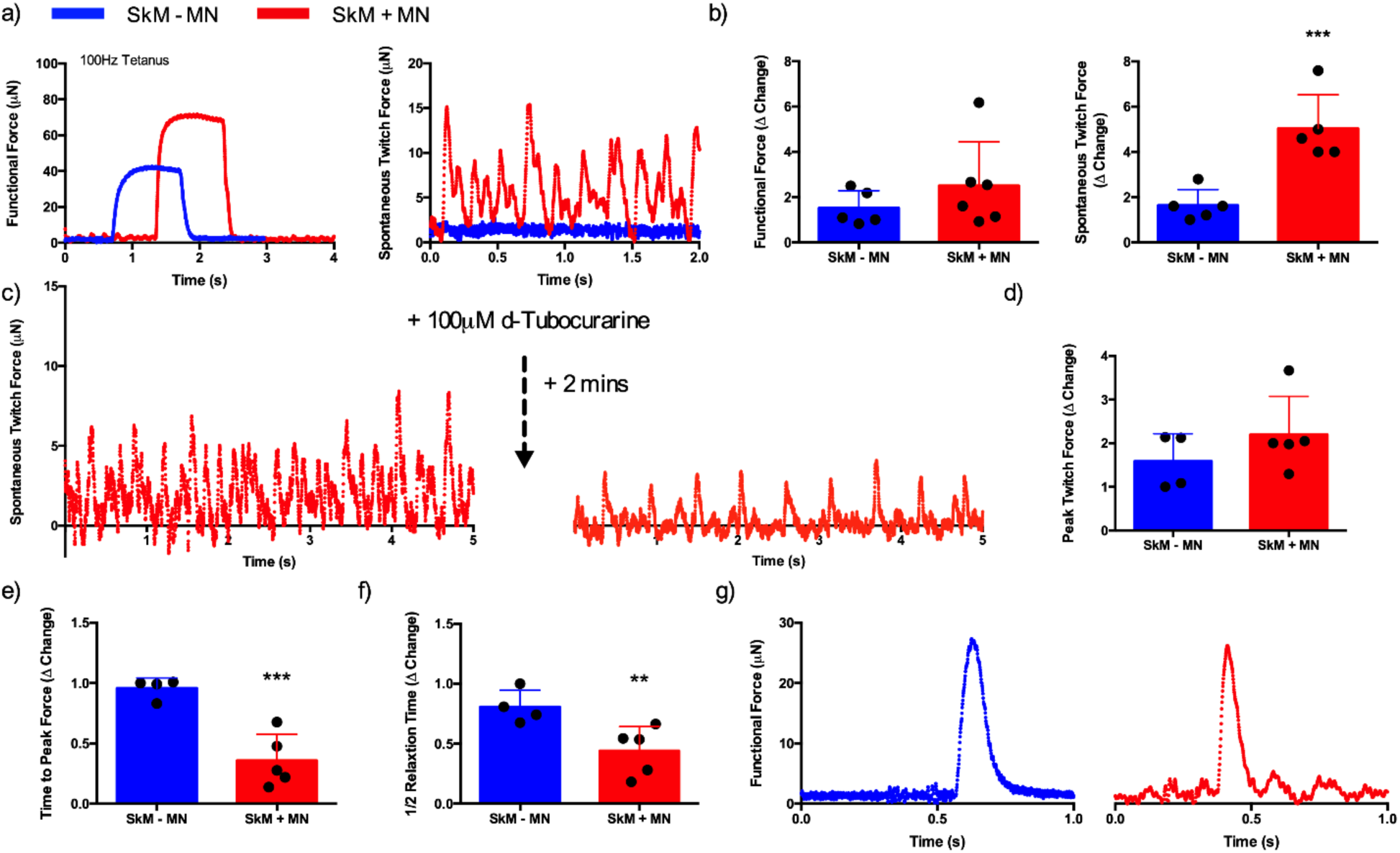
Physiological functionality of engineered tissues is enhanced via neuromuscular junction formation. (a) Maximal induced tetanus, and spontaneously occurring twitch force profiles in engineered tissues with (SkM +MN) and without (SkM -MN) iPSC motor neuron spheroids. (b) Quantification of induced tetanic and spontaneous twitch force driven via motor nerve inclusion. (c) Inhibition of the acetylcholine receptor within neuromuscular tissues via addition of 100µM d-Tubocurarine. (d) Maximal twitch force induced via electrical field stimulation. (e) Time to peak twitch force and (f) half relaxation time as physiological measures of contractile function. (g) Representative twitch profiles evidencing reduced time to peak twitch and shorter relaxation profiles in SkM + MN tissues. Δ change indicative of normalised data. Individual functional data points indicative of n=3 contraction profiles per tissue, totalling minimum of n=12 and maximum of n=18 contractions per condition and presented ± standard deviation (SD). **P ≤ 0.01 and ***P ≤ 0.001.

## 3. Discussion

New technologies are required that enable the study of the human NMJ in development, throughout disease progression and in response to therapeutic interventions. This work responds to this need, utilising a freely available 3D printed system to bioengineer human skeletal muscle and provides the methodologies to incorporate commercially available iPSC derived motor neuron progenitors to create functional human NMJs.^25^

iPSC motor neuron progenitors were cultured as 3D spheroids to enhance maturity and contain cellular soma separate from skeletal muscle tissues. Typical spheroid cultures utilise cell-cell adhesion to form multicellular aggregates that secrete extracellular matrix proteins.^28^ Enhanced transcription of motor neuron lineage markers in neural stem cell spheroids compared to monolayer cells has previously been reported.^29^ Our data supports this methodology, evidencing enhanced expression of developmental, progenitor, motor neuron and mature mRNAs, in addition to decreased proliferation of motor neuron progenitor spheroids compared to monolayer (Fig.1). Axon projections of cultured spheroids after 7, 14 and 21 days demonstrated developmental stage specific neurite lengths (Fig.2). Maximal axon extension was observed after 14 days spheroid culture, aligning with mRNA expression data. Transcription of Olig-2 and Islet-1 increase linearly until day 14 before reducing significantly by day 21, with only mature marker SMI32 further enhancing transcription at this time-point. This suggests that spheroids matured in suspension, as used in this work, require adherent surfaces prior to this point for optimal neurite growth. Differences in axon projections of neuronal stem cell and progenitor spheroids have been observed previously, with maximal axon lengths of <400μm being enhanced to ∼800μm following vascular integration.^29^ Importantly, this methodology to produce 3D spheroids results in axons, at 14 days, of 1000-1600µm length of which 90% are SMI-32+ (Fig.2) in the absence of a vascular component.

*In vivo* spinal motor neuron soma resides within a 3D environment of grey matter within the ventral horn, extending axons to the peripheral musculature. In an attempt to replicate this environment, different concentrations of type I collagen hydrogels were utilised to manipulate matrix properties. Concentration dependent increases in the transcription of genes indicative of maturity (SMI32), axon extension (MAPT) and vesicular transport of ACh (VACHT) neurotransmitter were evident in spinal motor neuron spheroids. This outlines that maturity and neurotransmitter function can be enhanced via addition to optimised 3D environments, and that the matrix composition, which differs from that required for skeletal muscle, affects expression of key mRNAs (Fig.3). Enhanced differentiation and axon growth of iPSC motor neuron spheroids has also been reported in soft methacrylated hyaluronic acid hydrogel compositions.^30^ However, previous work to load motor neuron spheroids to muscle tissue has used 2.4mg/mL concentrations of type I collagen and 2mg/mL supplemented with Matrigel™ in 4:1 ratios.^24,31^ This is contrasting to our data that demonstrates enhanced biological transcription at 1mg/mL, decreasing toward that of monolayer spheroids at lower (0.5mg/mL) and higher (1.7mg/mL) concentrations. This matrix concentration provides an optimised hydrogel for 3D culture of iPSC motor neuron spheroids and as a delivery vehicle to engineered muscle tissues using polymerised type I collagen. However, refining this matrix delivery system via inclusion of influential neural extracellular proteins may promote further enhancements in maturity and biological functionality for 3D motor neuron spheroid cultures.^32^

Multiple methods to bioengineer human skeletal muscle now exist using both primary and stem cell myogenic precursors.^25,33–35^ Consistent with previous research, 3D skeletal muscles in this work demonstrated enhanced number of AChR clustering across 3 week culture periods and representative pretzel-like morphologies compared to monolayer (Fig.4).^23^ This acceleration in development is mirrored in synaptic mRNAs for MuSK and Lrp4, which associate at the NMJ with z-agrin to form a protein complex across the synaptic basal lamina.^1,2^ Significant increases in MUSK mRNA were evident compared to monolayer throughout myogenic development, specifically notable after 2 days and 3 weeks culture. This may underpin the increased AchR clustering and morphological maturation observed in 3D tissues, as MuSK is known to regulate AchR clustering and organisation of the post-synaptic membrane in mice.^36,37^ This initial increase in transcription is also apparent in LRP4 mRNA, however, comparable levels are evident in monolayer myotubes after 1 week. At this point, monolayer expression has peaked and decreases, while mRNA levels in 3D tissue rise significantly until 2 weeks culture before also decreasing. Together, these data indicate upregulation of mRNA transcription for translation of the ‘core’ proteins expressed by the muscle required for NMJ formation and maintenance, and development of the post-synaptic apparatus for motor nerve synaptogenesis in 3D tissues after 2 weeks myogenic development.

Neuromuscular development in skeletal muscle is a multifaceted process of primary and secondary motor innervation that is highly dependent on myogenic status.^3^ Myogenic progenitors in the paraxial mesoderm either side of the neural tube proliferate and fuse to form immature primary myotubes.^38^ Primary myotube developmental stages precedes that of axon penetration from the cervical and brachial plexus in the diaphragm of mice and rats.^39,40^ Following primary innervation, residual fetal myoblasts fuse and generate secondary myotubes.^38^ However, the formation of secondary fibres requires the presence of the motor nerve, which influences post-synaptic specialisation such as contractile properties of fibres.^41^ Previous work has introduced the motor nerve at experimental onset prior to primary myotube formation and following initial myogenesis, both generating functional neuromuscular tissues.^23,24^ In our model, attempts to incorporate motor neuron spheroids with HDMCs at experimental onset did not form functional tissues. However, this is likely due to the requirement for complex medium formulations that simultaneously support differentiation phases of both cell types, opposed to developmental issues (Fig.S4). Engineered tissues in this work fuse to form functional primary myotubes after 14 days culture in the absence of motor axons. At this point, motor neuron addition enables primary innervation of myotubes with a further 2 week culture period to allow for secondary fibre formation and innervation. The paucity of data regarding human NMJ formation provides difficulties when devising methodologies that accurately represent neuromuscular development *in vitro*. However, systems that enable the precise addition of motor neurons dependent on myogenic status, as reported in this work, likely afford the greatest utility in this endeavour.

Motor recruitment of skeletal muscle fibres *in vivo* is performed in a size dependent manner by *α* motor neurons.^42^ Typically, singular motor neurons regulate recruitment of multiple groups of fibres via axonal branching comprising one motor unit, with multiple motor units necessary to maximally contract entire muscles. To investigate the motor pool required to maximally contract engineered muscle tissues in this work, motor neuron spheroids were loaded, following 2 weeks maturation (muscle and motor nerve), at ratios of 2, 4 or 6 per tissue. Negative linear relationships in spontaneous muscle contractions were observed indicating a maximal capacity for spheroid addition in this model (Fig.5). Regarding previous human NMJ models, monolayer cell clusters were dosed to engineered muscle tissues at a ratio of 3 per ∼10mm tissue.^23^ Our data aligns with these ratios, indicating maximal fibre recruitment when loading 1 motor nerve per ∼3mm of engineered muscle tissue length. Importantly, these data also evidence a threshold of 1 spheroid per ∼1mm of muscle tissue length at which innervation and or motor recruitment appears inhibited. This is contrasting to that observed in the microfluidic platform to create neuromuscular tissues, that loaded a singular motor spheroid per muscle tissue of approximately 1.5mm lengths.^24^ This may allude to competition for nutrient and or growth factor availabilities as the major determinant. Motor neuron spheroids are matured individually in 200μL volumes, with motor unit tissues residing in 2mL of supplemented medium. This would provide adequate growth factor concentrations even at highest spheroid ratios. However, given the introduction of the muscle tissue that has also been known to utilise neuronal growth factors such as BDNF in mice, further research is required to establish the biochemical or physical causality of this effect.^43^

High levels of pre- and post-synaptic co-localisation with adjacent neuronal axon terminals were apparent in neuromuscular tissues with ∼13 synaptic contacts evident within a singular z-slice, equating to 4.6 NMJs per mm^2^. High magnification images of these contacts showed features consistent with human NMJ morphologies, in addition to motor neve axons tracking along myosin heavy chain positive fibres (Fig.6). Human NMJ models must enable morphological and functional characterisation to provide complete utility in physiological and or pathophysiological studies, in addition to sensitive protein and RNA analyses. Previous studies have provided functional data, yet limited morphological characterisation other than adjacent Tuj-1 positive axons and muscle fibres.^24,31^ The most sophisticated morphological characterisation of 3D tissues to date identified core synaptic proteins of laminin-*β*2, MuSK and rapsyn at the motor nerve-muscle interface.^23^ Although thorough functional analysis was also performed, all aforementioned proteins are either expressed or accumulated by the post- synaptic membrane.^3^ To ensure full biochemical characterisation it is recommended to fluorescently label proteins of both pre- and post-synaptic functionality (AChR and synaptic vesicle markers; synaptotagmin, SV- 2) as typically utilised within *in vivo* models and evidenced within this work.

Physiological assessments of motor unit function confirmed spontaneous contraction profiles were driven via the NMJ with inhibition of the post-synaptic membrane using AChR antagonists. This method has widely been utilised amongst the literature *in vitro*.^14,22^ Notably, no significant differences in induced maximal tetanic or twitch contraction forces were evident in innervated tissues. This aligns with microfluidic human 3D neuromuscular co-cultures, but contrasts with those produced using primary rat derived myogenic and motor neuron cells.^22,24^ This work is, however, the first to evidence that NMJ formation within bioengineered human motor units enhances the neuromuscular physiology measures typically used to assess neuromuscular function *in vivo*.^44^ Following release of ACh and binding to the AChR, the resulting depolarisation opens volatage gated sodium ion channels creating an action potential that propagates along the cell, invades the t-tubules and elicits opening of L-type calcium channels. Ryanodine receptors (RyRs) in the sarcoplasmic reticulum (SR) then open and release calcium required for muscular contraction. Reductions in cytosolic calcium is then achieved via SR re-uptake mediated the sarco-endoplasmic reticulum calcium ATPase (SERCA) pump eliciting muscle relaxation.^4^ Time to peak twitch is typically indicative of skeletal muscle excitability, which is underpinned by the efficiency of neuromuscular transmission, propagation of action potential and release of calcium via the SR RyRs. Whereas decreased half relaxation time has been outlined to be indicative of calcium re-uptake in the SR via the SERCA pump, with increased SERCA1 mRNA transcription evident following electrical stimulation of 3D muscle tissue.^45^ Both functional measures were significantly decreased in time following innervation, indicating an enhanced efficiency of the molecular mechanisms of muscle contraction. Importantly, neuromuscular tissues using primary rat cells reported increases in time to peak force and half relaxation time measures upon innervation, suggesting that alterations in the excitability of bioengineered motor units appears sensitive across species. This further outlines the importance of models derived entirely from human cellular material for the study of cell/molecular biology and physiology of the human neuromuscular junction.

In summary, this work presents the methods required to fully maximise maturity of both iPSC motor neurons and primary skeletal muscle, utilising cell type specific extracellular matrices and developmental timelines for the study of human neuromuscular physiology. Future work regarding this model should seek to fully characterise the developmental stages of NMJ formation and function in humans. Focus should also be applied on the integration of additional neuronal cell types such as non-myelinating and myelinating schwann cells, and sensory neurons to complete the neuromuscular reflex-arc. Finally, this model should be translated to provide a functional assay of NMJ pathophysiology via integration of patient iPSC motor neurons.

## 4. Materials and methods

### 4.1. Primary human derived myoblast cell purification and culture

This study was approved by the Loughborough University Ethics Approvals (Human Participants) Sub Committee (reference number: R18-P098). Healthy male subjects provided written informed consent and completed a medical screening questionnaire prior to participation. Primary human derived myoblast cell (HDMC) samples were obtained from healthy males (n = 3), between the ages of 18–55 reporting no recent injuries or intake of anti-inflammatory pharmaceuticals, from the vastus lateralis muscle using routine muscle biopsy procedure with micro-biopsy. HDMCs were isolated using an established explant protocol and sorted for the expression of CD56, with detailed methods of the biopsy procedure and cell purification methodology being available in the Supplementary Information. Upon resuscitation, HDMC CD56+ and - cells were expanded separately until passage 7/8 for use then remixed at a ratio of 9:1 (+/-). Remixed HDMCs were plated for experimental use in monolayer at 1×10^4^ cells/cm^2^ and cultured in growth medium (GM); composed of 79% Dulbecco’s modified Eagles medium (DMEM; Sigma), 20% fetal bovine serum (FBS, Pan Biotech) and 1% penicillin− streptomycin (P/S, Fisher) until confluence, prior to inducing differentiation via addition of differentiation medium (DM) composed of; 97% DMEM, 2% horse serum (Sigma), 1% P/S supplemented with 20ng/mL insulin-like growth factor 1 (IGF-1, PeproTech). HDMC medium was replenished entirely at 48h intervals.

### 4.2. iPSC derived motor neuron monolayer and spheroid culture

Following resuscitation, iPSC derived motor neuron progenitors (Axol Bioscience, UK, ax0078) were seeded at 2×10^6^ cells/T25 culture flask coated with 1.5mg/mL Corning™ Matrigel™ (Fisher) DMEM solution, in motor neuron recovery medium (RM, Axol) supplemented with 0.1µM all-trans retinoic acid (RA, Sigma) and 10µM Y-27332 ROCK inhibitor (Stem Cell Technologies, removed after 24h). At cellular confluence, motor neuron progenitors were dissociated using TrypLE select (1x, Gibco, Fisher), centrifuged at 200 G for 5 minutes and seeded in 96-well U bottom spheroid microplates (MoBiTec, MS-9096UZ) at 1×10^4^ cells/well or monolayer at 1×10^5^ cells/cm^2^ in motor neuron maintenance medium (MM, Axol) supplemented with 0.5µM RA, 5ng/mL brain derived neurotrophic factor (BDNF, PeproTech) and 10ng/mL ciliary neurotrophic factor (CNTF, Axol). Monolayer motor neuron progenitors MM for cell seeding was supplemented with 10µM Y- 27332 ROCK inhibitor to promote adhesion and survival. All medium for both monolayer and 3D spheroids were changed completely after 24h and then replenished by 50% (half-change) thereafter at 48h intervals. Monolayer and spheroid motor neurons were isolated for mRNA analyses at 2, 7, 14 and 21 days of maturation. For analysis of motor neuron spheroid axonal extension at varying developmental stages (7, 14 and 21 days), spheroids were removed from microplate wells, individually seeded in complete MM on Matrigel™ coated (1.5mg/mL) well plates and left to extend neurons for 5 days prior to fixation.

### 4.3. 3D skeletal muscle and motor neuron tissue engineering

Skeletal muscle tissue engineering was undertaken as previously reported via addition of 65% v/v type I rat tail collagen; dissolved in 0.1M acetic acid, protein at 2.035mg/mL (First Link, UK), with 20% Matrigel™ matrix and 10% v/v of 10× MEM (Fisher).^25^ This solution was neutralized by the dropwise addition of 5 and 1M sodium hydroxide (NaOH, Sigma), until a colour change to cirrus pink was observed. HDMC cells were added at a seeding density of 4×10^6^ cells/mL in a 5% v/v GM solution to 3D printed moulds and incubated for 10−15 minutes (37°C, 5% CO_2_). Engineered skeletal muscles were then maintained in 2mL GM and cultured for a further 14 days (4 days GM, 10 days DM). To optimise the preferred neuronal type I collagen matrix concentration, iPSC motor neuron spheroids were removed from microplates and added to a pre-neutralised stock collagen/MEM hydrogel solution (1.73mg/mL) further diluted with MM to yield the following concentrations: 1.7mg/mL, 1mg/mL and 0.5mg/mL at 500µL volumes in culture well plates. Following neutralisation, each 3D motor neuron hydrogel and was maintained in MM and cultured alongside monolayer spheroid conditions, prepared as previously described, and cultured for 5 days prior to isolation for genetic and morphological analyses. Spheroids were also cultured in monolayer on 1.7mg/mL, 1mg/mL and 0.5mg/mL coated surfaces, however this reduced adherence and axonal extension compared to Matrigel™ coated plates 369(Fig.S3).

### 4.4. 3D neuromuscular tissue engineering

Engineered primary HDMC tissues are cultured for 14 days as described above, prior to being encased within an additional extracellular matrix, comparable in hierarchy to the perimysium, that contains iPSC derived motor neuron spheroids. The maturity of the motor neuron spheroids is standardised at the optimised time- point (14 days) for axonal extension and maturation as is the co-culture time-period (14 days). To combine motor neuron spheroids into appropriate hydrogel volumes/concentrations, spheroids are extracted and placed at desired numbers in sterile Eppendorf tubes®, centrifuged at 100 G for 30s, with the supernatant discarded prior to resuspension within desired loading matrix (Fig.S1). Spheroids are then added around pre-differentiated HDMC tissues (day 14) within an optimised type I collagen (1mg/mL) perimysium matrix and cultured for a further 14 days within MM (Fig.S2).

### 4.5. Functional assessment of skeletal muscle and neuromuscular tissues

Engineered skeletal muscle and neuromuscular tissues were washed twice in phosphate buffered saline (PBS) prior to one end of the engineered muscle being lifted from the pseudo tendon pin. The loose end of the construct was then attached to the force transducer (403A Aurora force transducer, Aurora Scientific) using the eyelet present in the construct. For direct electrical stimulation, the tissue was submerged (3mL) in Krebs-Ringer-HEPES buffer solution (KRH; 10mM HEPES, 138mM NaCl, 4.7mM KCl, 1.25mM CaCl_2_, 1.25mM MgSO_4_, 5mM Glucose, 0.05% Bovine Serum Albumin in dH_2_O). Wire electrodes were positioned parallel either side of the construct to allow for electric field stimulation. Impulses were generated using LabVIEWsoftware (National Instruments, UK) connected to a custom-built amplifier. Maximal twitch force was determined using a single 3.6V/mm, 1.2ms impulse and maximal tetanic force was measured using a 1s pulse train at 100Hz and 3.6V/mm, generated using LabVIEW 2012 software (National Instruments, UK). For measurement of spontaneous muscle contraction driven by the motor nerve in neuromuscular tissues, once attached, each tissue sample was recorded for a period of ≥ 30s prior to further analysis. Antagonistic blocking of the AChR was achieved via 1mL bolus of 200µM d-tubocurarine (Sigma) to 1mL of surrounding basal medium (MM), achieving 100µM concentration as previously reported.^22^ Where possible induced twitch and tetanus data were derived from 3 contractions per tissue sample. Data was acquired and analysed using a Powerlab system (ver. 8/35) and associated software (Labchart 8, AD Instruments, UK).

### 4.6. Immunocytochemistry and morphological analysis

To perform immunocytochemistry at experimental termination, monolayer and 3D samples were washed twice in 2mL of PBS/well and fixed using a 3.7% paraformaldehyde solution (Sigma). Monolayer and 3D samples were then permeabilised (0.2% Triton X-100, Fisher) and blocked using 5% goat serum (Fisher) for 30 minutes, prior to being incubated overnight (≥12h) with primary antibody solutions. Primary antibodies were co-incubated with 5% goat serum to reduce non-specific binding. Labelled samples were then counterstained with secondary antibody fluorophores and fluorescent small molecules for ≥2h (Table S1). Samples were washed 3 times (1× tris-buffered saline (TBS)) before permeabilisation and after both primary and secondary antibody incubations.

Fluorescence images were captured using a Leica DM2500 fluorescence microscope with manufacturer’s software (Leica Application Suite X). Confocal imaging was undertaken on a Zeiss LSM 880, using 40x and 63x oil objectives. Images captured via confocal tile scan are stitched together enabling visualization of large tissue sections. One standard scan equates to a 3 × 7 tile image, taken in a single z-plane from the centre of the engineered tissue. Scans used to identify synaptic contacts (AChR:SV-2:Tuj-1 and AChR:Tuj-1:MF-20) and co-localisation in neuromuscular tissues are 10 × 8 and 7 × 5 stitched visualisations respectively. Images were analysed using IMAGE J 1.50a/Fiji (Java 1.6.0_24) software (National institute of Health, USA).^26^ Monolayer image inclusion criteria (n = 1) were set at ≥5 images taken at random locations per well. Myotube inclusion criteria were defined as containing ≥3 nuclei per myotube; 2 nuclei constituting a dividing cell and 3 or more indicating myogenic fusion. Myotube width analysis was performed via measuring the central and representative region of each myotube within an image. Analysis of AChR size and number was performed using an in-house macro designed for Fiji (Java 1.6.0_24) image analysis software (Image J 1.50a). Analysis of motor neuron axonal extension in spheroids at 7, 14, and 21 days maturation was undertaken using a coordinate analysis system. Coordinates are assigned to the centre of the spheroid and to all adjacent axon terminations. Axon termination X and Y coordinates are then subtracted from respective central equivalents to yield axon length (µm).

### 4.7. RNA extraction and quantitative real-time polymerase chain reaction

Tissue-engineered skeletal muscle was thawed and suspended in 500μL TRIzol™ (Fisher) before being homogenized using TissueLyserII (Qiagen) for ≥2 minutes until tissue degradation was complete. Monolayer and motor neuron spheroid RNA was extracted using the TRIzol™ method, according to manufacturer’s instructions (Sigma). RNA concentration and purity were obtained by UV-Vis spectroscopy at optical density of 260 and 280nm using a Nanodrop 2000 (ThermoFisher Scientific). All RNA samples were analysed in duplicate. Five nanograms of RNA were used per real-time polymerase chain reaction (RT-PCR) for RPIIβ, PAX6, OLIG2, ISLET1, SMI32, MKI67, MAPT, VACHT, MYOD, MYOG, LRP4 and MUSK (Table S2). RT-PCR amplifications were carried out using a Power SYBR Green RNA-to-CT 1 step kit (Qiagen) on a ViiATM Real-Time PCR System (Applied Biosystems, Life Technologies), analysed using ViiATM 7RUO software. The RT-PCR procedure was as follows: 50°C, 10 minutes (for cDNA synthesis), 95°C, 5 minutes (transcriptase inactivation), followed by 95°C, 10s (denaturation), 60°C, 30s (annealing/extension) for 40 cycles. Relative gene expression was calculated using the comparative C_T_ (ΔΔC_T_) equation for normalized expression ratios; relative expression calculated as 2 − ΔΔC_T_, where C_T_ is representative of the threshold cycle.^27^ RPII-β was used as the housekeeping gene in all RT-PCR assays. To compare conditions, one control sample from each experimental repeat (n =3) was used as the calibrator condition in the C_T_ (ΔΔC_T_) equation. RT- PCR data is presented as the relative gene expression level, determined by the ΔΔC_T_ equation.

### 4.8. Statistical analyses

Significance of data were determined using IBM© SPSS© Statistics version 23. Mauchly’s test of sphericity and Shapiro-Wilk tests were used to confirm homogeneity of variance and normal distribution of data respectively. Where parametric assumptions were met, factorial analysis of variance (ANOVA) was performed with Bonferroni post hoc analyses used to analyse differences between conditions at specific time-points. Nonparametric Kruskal−Wallis (H) analysis was undertaken where data violated parametric assumptions. Mann Whitney (U) tests were then utilised to determine the significance between conditions, in accordance with Bonferroni correction to account for incremental type-1 error. All data is reported as mean ± standard deviation (SD). Statistical significance was assumed at P ≤ 0.05.

## Acknowledgements

Authors would like to thank the Rosetrees Trust and Stoneygate Trust (Grant ref: M889), Axol Bioscience, and Loughborough University for supporting this work.

