## Supplementary Information for "Bioengineered model of the human motor unit with physiologically functional neuromuscular junctions"

Supplementary Figures
Fig.S1 – Extraction and centrifugation of iPSC motor neuron spheroids.
Fig.S2 – Addition of acellular type 1 collagen matrix to tissue engineered skeletal muscle does not affect morphology or force production.
Fig.S3 – Matrix surface coating dictates iPSC derived motor neuron spheroid axonal extension.
Fig.S4 – Co-culture at experimental onset requires complex medium formulations that support differentiation of both myogenic and motor neuron cells.
Fig.S5 – Stitched fluorescence microscopy visualisation of neuromuscular tissues.

Supplementary Tables
Table S1 – Antibodies and chemical/secondary fluorochromes utilised for the identification target proteins.
Table S2 – Primers utilised for the examination of target mRNA.

1. SI Text
1.1. Primary human derived myoblast cell purification and culture

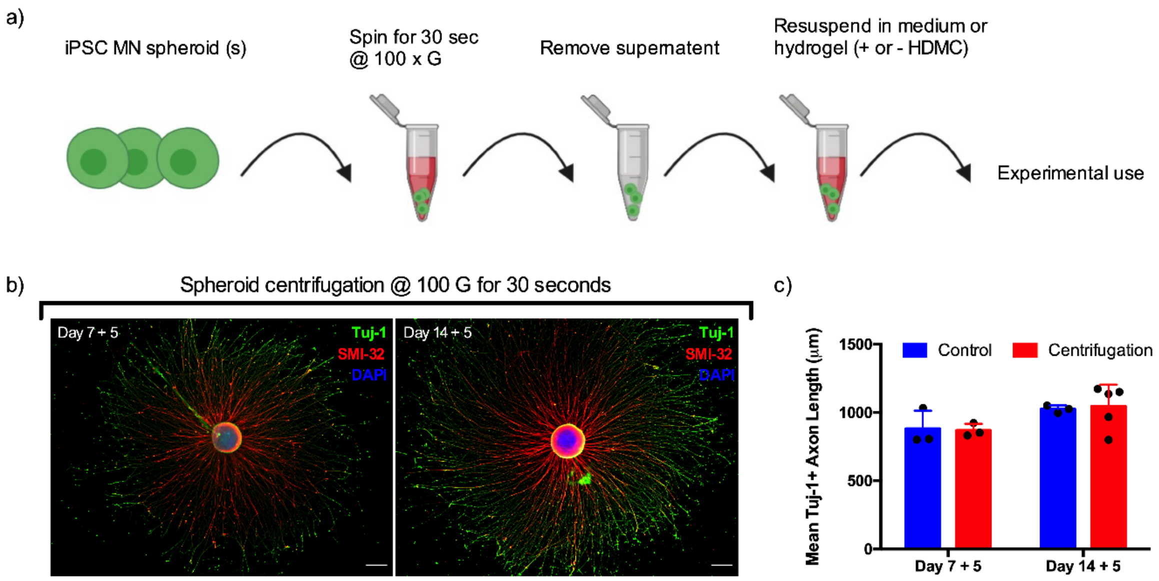


Fig.S1. **Extraction and centrifugation of iPSC motor neuron spheroids**. (a) Schematic detailing process for extraction and resuspension of iPSC derived motor neuron spheroids for experimental use. (b) Fluorescently labelled spheroids for Tuj-1 and SMI-32 following extraction at 7 or 14 days and cultured for a subsequent 5 days. (c) Quantification of spheroid axons evidencing no effect of centrifugation at 100xG for 30 seconds. Scale bars = 200 µm.

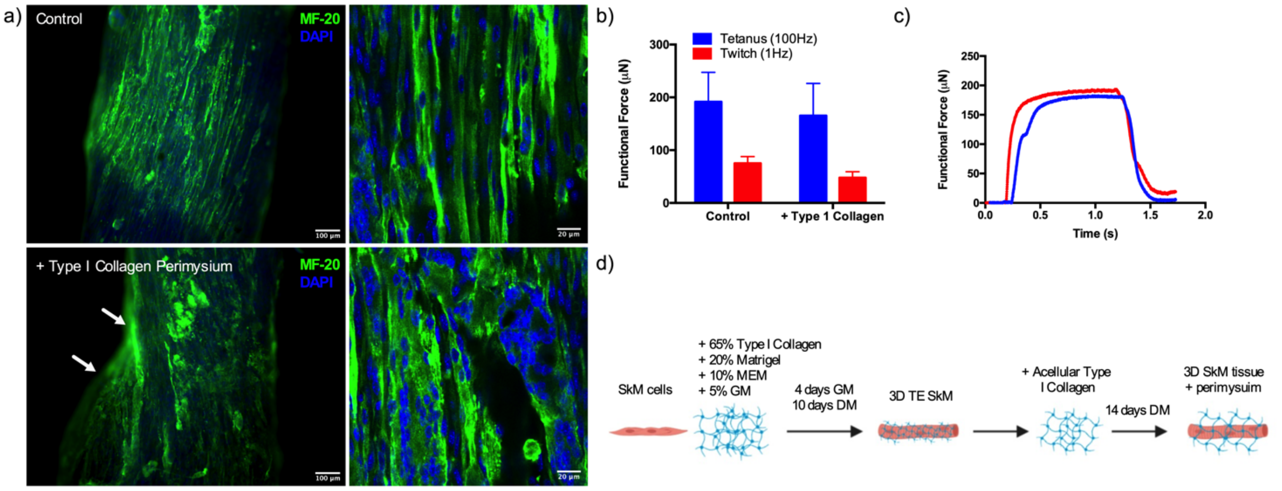


Fig.S2. **Addition of acellular type 1 collagen matrix to tissue engineered murine skeletal muscle does not affect morphology or force production.** (a) Engineered C2C12 skeletal muscle tissues labelled for myosin heavy chain (MF-20) cultured for 4 weeks without (control) or following addition of acellular type I collagen perimysium (1mg/mL) for the final 2 weeks of culture. (b) Tissue functionality assessed via electrical field stimulation for maximal tetanic and twitch contractions. (c) Representative tetanus force traces with and without type I collagen sheath. (d) Schematic detailing method for addition of acellular matrix. Scale bars = 100 µm (left) and 20 µm (right).

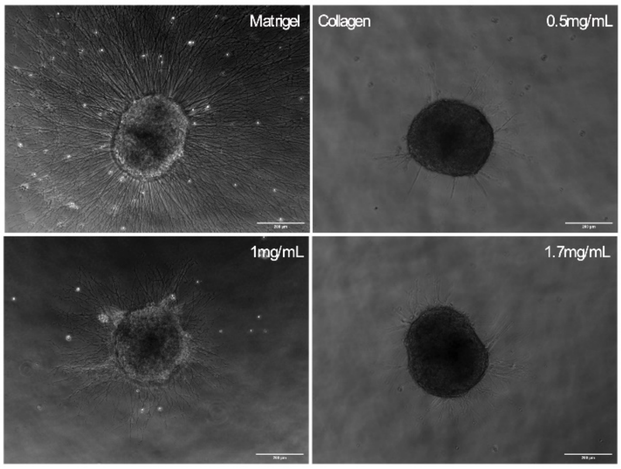


Fig.S3. **Matrix surface coating dictates iPSC derived motor neuron spheroid axonal extension.** iPSC MN spheroids were cultured for 14 days to optimised developmental stage prior to being adhered to either Matrigel, or collagen at concentrations of 0.5, 1 and 1.7mg/mL. Following 5 days culture, all MNs adhered to the substrates, however, the outgrowth of axons was significantly reduced in all collagen concentrations. This data was undertaken in parallel with that displayed in Fig.3. Together, these data indicate that providing a 3D environment to MN spheroids enhances the transcription of mature MN genes, and that this is likely related to the stiffness of the substrate opposed to the molecular composition or concentration of proteins for adherence. Scale bars = 200µm.

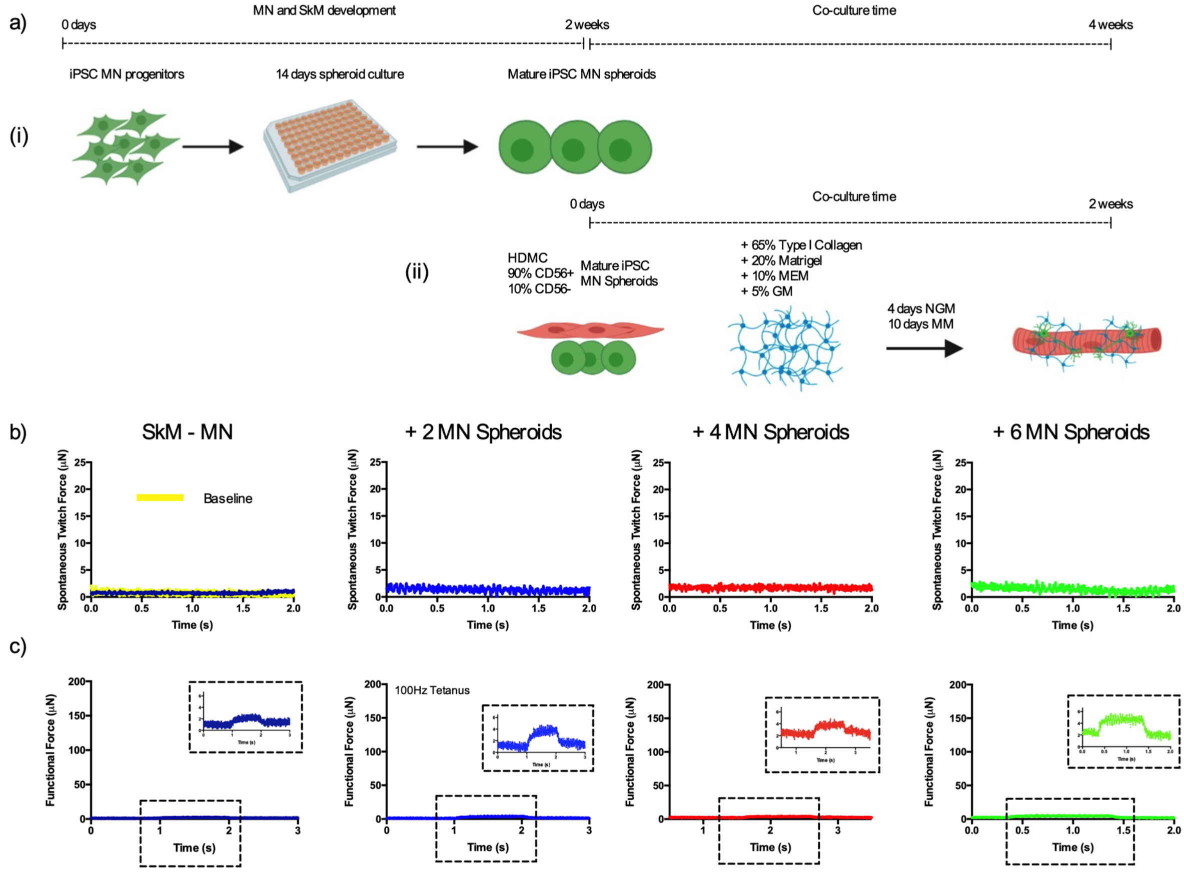


Fig.S4. **Co-culture at experimental onset requires complex medium formulations that support differentiation of both myogenic and motor neuron cells.** (a) Schematic detailing method for inclusion of SkM and MN progenitors at experimental onset. (i) MN progenitors are loaded to spheroid microplates and matured for 14 days. (ii) Donor matched HDMCs are added to established SkM hydrogel simultaneously with matured MNs and cultured for 2 weeks. (b) Seeding of 2, 4 or 6 motor neuron spheroids with skeletal muscle cells do not exhibit increases in functional force or (c) spontaneous twitch indicative of motor nerve induced contraction. NGM composed of; 79% DMEM/F12, 20% FBS, 1% P/S, supplemented with 0.5µM RA, 5ng/mL BDNF, 10ng/mL CNTF and 100µM ascorbic acid. Abbreviations: MN; Motor neuron, SkM; Skeletal muscle, HDMC; Human derived muscle cells, GM; Growth medium, DM; Differentiation medium, MEM; 10X Minimum essential medium, NGM; Neuromuscular growth medium, MM; Motor neuron maintenance medium.

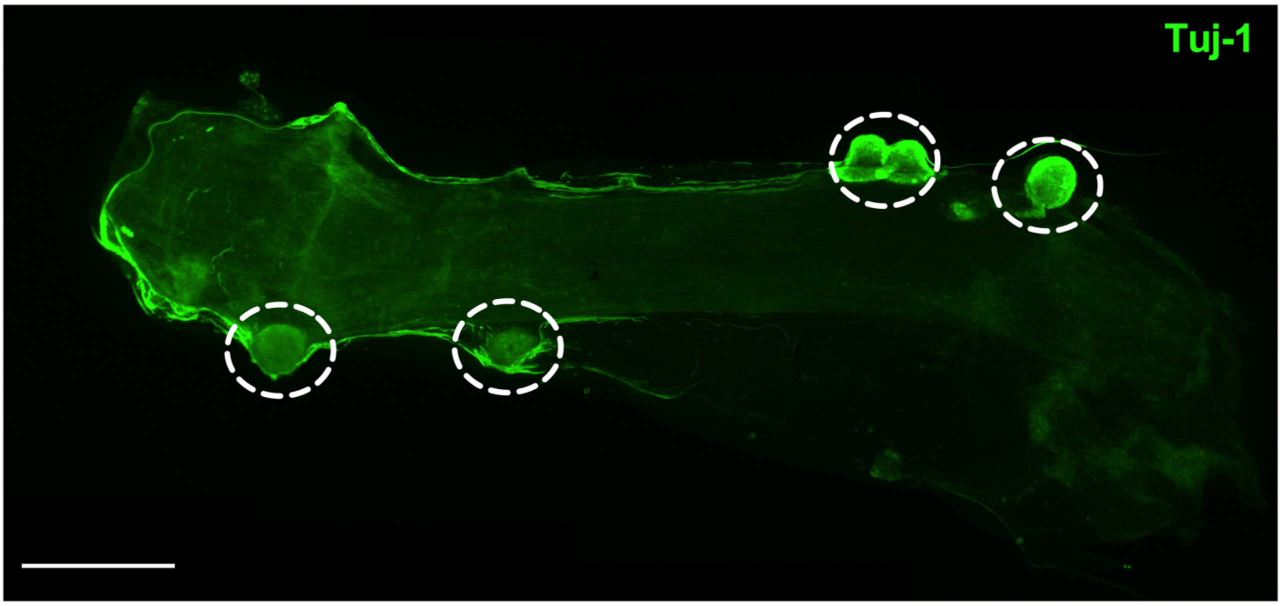


Fig.S5. **Stitched fluorescence microscopy visualisation of neuromuscular tissues.** Neuromuscular tissues loaded with 4 motor neuron spheroids positive for neuronal cytoskeletal marker Tuj-1 following 2-weeks co-culture (indicated via white circular surrounds). Following addition of iPSC motor neurons within collagen hydrogels, CD56- myogenic cells migrate into and remodel loaded matrix to pull motor neuron spheroids tight to skeletal muscle tissues. Scale bar = 1mm.

**Table S1** – Antibodies and chemical/secondary fluorochromes utilised for the identification of target proteins.

| Target | Antibody | Species | Dilution | Distributor |
|---|---|---|---|---|
| Olig-2 | Recombinant Anti-Olig2 antibody [EPR2673] (ab109186) | Rabbit | 1:500 | Abcam |
| HB-9 | Anti-HB9 antibody (ab221884) | Rabbit | 1:500 | Abcam |
| SMI-32 | Anti-Neurofilament heavy polypeptide antibody (ab8135) | Rabbit | 1:200 | Abcam |
| Tuj-1 | Recombinant Anti-beta III Tubulin antibody [EP1569Y] (ab52623) | Rabbit | 1:500 | Abcam |
| N/A | Goat Anti-Rabbit IgG H&L (Alexa Fluor® 488) (ab150077) | Goat | 1:500 | Abcam |
| N/A | Goat Anti-Mouse IgG H&L (Alexa Fluor® 488) (ab150113) | Goat | 1:500 | Abcam |
| N/A | Goat Anti-Rabbit IgG H&L (Alexa Fluor® 647) (ab150079) | Goat | 1:500 | Abcam |
| N/A | Goat Anti-Mouse IgG H&L (Alexa Fluor® 647) (ab150115) | Goat | 1:500 | Abcam |
| AChR | Invitrogen™ Molecular Probes™ α-Bungarotoxin, Alexa Fluor™ 594 Conjugate | N/A | 1:500 | Fisher |
| | Invitrogen™ Molecular Probes™ α-Bungarotoxin, Alexa Fluor™ 647 Conjugate | N/A | 1:500 | Fisher |
| Pan-Myosin heavy chain | MF20 monoclonal - MF 20 was deposited to the DSHB by Fischman, D.A. (DSHB Hybridoma Product MF 20) | Mouse | 1:200 | DSHB |
| Nuclear DNA | DAPI (4',6-diamidino-2-phenylindole, dihydrochloride) | N/A | 1:2000 | Fisher |
| SV-2 | SV2 monoclonal - SV2 was deposited to the DSHB by Buckley, K.M. (DSHB Hybridoma Product SV2) | Mouse | 1:20 | DSHB |

**Table S2** – Primers utilised for the examination of target mRNA.

| Target mRNA | Primer Sequence (5'-3') | Product Length | NCBI Reference Sequence |
|---|---|---|---|
| RPII-β | F: AAGGCTTGGTTAGACAACAG<br>R: TATCGTGGCGGTTCTTCA | 142 | NM_000938.3 |
| MYOD | F: ACGGCATGATGGACTACAGC<br>R: TGGCAGTCTAGGCTCGACAC | 132 | NM_002478.5 |
| MYOG | F: CAGCTCCCTCAACCAGGAG<br>R: GCTGTGAGAGCTGCATTCG | 91 | NM_002479.6 |
| MUSK | F: AGACAGCCCTCTCAGGGAAA<br>R: CCTGGCAAAAACCTCAACTTCC | 169 | NM_005592.4 |
| LRP4 | F: AAATCCAAGTTCACTGATCC<br>R: CCTCTTTCTTATAGCACAGC | 136 | NM_002334.4 |
| PAX6 | F: AGTGCCCGTCCATCTTTGC<br>R: CGCTTGGTATGTTATCGTTGGT | 81 | NM_001127612.2 |
| OLIG2 | F: CCAGAGCCCGATGACCTTTTT<br>R: CACTGCCTCCTAGCTTGTCC | 178 | NM_005806.4 |
| ISLET1 | F: GCGGAGTGTAATCAGTATTTGGA<br>R: GCATTTGATCCCGTACAACCT | 102 | NM_002202.3 |
| SMI32 | F: GCAGTCCGAGGAGTGGTTC<br>R: CGCATAGCGTCTGTGTTCA | 75 | NM_021076.4 |
| MKI67 | F: ACGCCTGGTTACTATCAAAAGG<br>R: CAGACCCATTTACTTGTGTTGGA | 209 | NM_001145966.2 |
| MAPT | F: GAAAAATAGGCCTTGCCTTAG<br>R: CCTTGAGTTTCATCTCCTTTG | 170 | XM_005257371.4 |
| VACHT | F: CTGCTAGTGAACCCCTTGAGC<br>R: CAGGACTGTAGAGGCGAACAT | 99 | NM_003055.3 |

### 1. SI Text

#### 1.1. Primary human derived myoblast cell purification and culture

Tissue was obtained via the micro-biopsy procedure, with any visible connective tissue being removed. Tissue samples were removed from the storage GM solution and washed three times in a buffer solution (phosphate-buffered saline (PBS), 1% P/S & 1% Amphotericin, Sigma, UK). Once washed tissue chunks were placed into a petri dish, suspended in 1mL of growth medium (GM); composed of 79% Dulbecco's modified Eagle medium (DMEM; Sigma), 20% fetal bovine serum (FBS, Pan Biotech) and 1% penicillin− streptomycin (P/S, Fisher), and mechanically minced using 2 scalpel blades until broken down into small sized pieces. Tissue was then seeded into 0.2% gelatin (v/v PBS, Fisher) coated T25 flasks (approximately 4 pieces/flask) and suspended in 0.5mL of GM to ensure tissue was planted on the culture surface and not floating. Flasks were then placed in standard tissue culture incubators (37ºC humidified atmosphere/5% $CO_2$) for 7–10 days to allow cellular migration to occur with more GM added to prevent flasks drying out. Migration of HDMC's was monitored with the migrated cellular population passaged at 60% confluence to prevent spontaneous differentiation at low passage.

To purify the myogenic population at passage 3, cells were dissociated with accutase (Fisher-Scientific, UK) with the resulting pellet resuspended in 1mL of sterile filtered MACS© running buffer (PBS, pH 7.2, 0.5% bovine serum albumin (BSA), and 2 mM EDTA). A pre-separation filter (Miltenyi Biotech, UK) was prepared by washing once with 500µL running buffer, then the cells passed through in two 500µL batches. Following cells, three 1mL washes with running buffer were performed. All filter run through was collected excepting the initial preparatory wash. The resulting cell suspension was then centrifuged (300 x G for 10 mins) and the pellet resuspended in 80µL running buffer and 20µL of CD56 magnetic microbeads (Miltenyi). This cell suspension was kept dark and incubated (15 mins, 4°C) with gentle mixing every five minutes. Following incubation, the cell/bead suspension was centrifuged (300 x G for 10 mins) and the pellet resuspended in 500µL of running buffer. One midiMACS© column was mounted on the separator and prepared by passing 2mL of running buffer through the dry column, any runoff was discarded. Unsorted cells were applied to the top of the column and washed through (3 x 3mL) with running buffer. The effluent of the column was collected and represented the CD56- population. The column was then removed from the separator and 5mL of running buffer added, following buffer addition the column was plunged and the resulting effluent collected containing the CD56+ population. CD56- cells were plated on 0.2% gelatin coated flasks as described above. CD56+ cells were plated in flasks coated with Matrigel (1.5mg/mL in DMEM). Both CD56- and + fractions are then expanded until passage 5 and cryopreserved for experimental use.
